# Changes in Anterior and Posterior Hippocampus Differentially Predict Item-Space, Item-Time, and Item-Item Memory Improvement

**DOI:** 10.1101/551705

**Authors:** Joshua K. Lee, Yana Fandakova, Elliott G. Johnson, Neal J. Cohen, Silvia A. Bunge, Simona Ghetti

**Author notes:** Corresponding Authors: Joshua K Lee; Simona Ghetti.

## Abstract

Relational memory requires the hippocampus, but whether distinct hippocampal mechanisms along the anterior-posterior axis are required for different types of relations is debated. We investigated the contribution of structural changes in hippocampal head, body, and tail subregions to the capacity to remember item-space, item-time, and item-item relations. Memory for each relation and volumes of hippocampal subregions were assessed longitudinally in 171 participants across 3 time points (*M*_*age*_ at T1= 9.45 years; *M*_*age*_ at T2= 10.86 years, *M*_*age*_ at T3=12.12 years; comprising 393 behavioral assessments and 362 structural scans). Among older children, volumetric growth in: (a) head and body predicted improvements in item-time memory, (b) head predicted improvements in item-item memory; and (c) right tail predicted improvements in item-space memory. The present research establishes that volumetric changes in hippocampal subregions differentially predict changes in different aspects of relational memory, underscoring a division of labor along the hippocampal anterior-posterior axis.

---

Without the ability to retain relational information about life events our memories would be fragmentary, difficult to retrieve, and ultimately of little value. Relational memory depends on mechanisms that bind features of experiences into integrated event representations ^1^; these features include where an event happened (item-space)^2^, when it happened (item-time)^3^, and with what other events it co-occurred (item-item)^4^. The hippocampus is critical for learning and recalling these arbitrary memory relations ^5,6^, but whether all types of memory relations are supported by the same or different hippocampal mechanisms is debated ^7–9^.

On the one hand, there is substantial evidence that the hippocampus is necessary to learn all arbitrary relations. For example, Konkel and colleagues found that adults with hippocampal lesions were equally impaired in their ability to remember spatial, temporal, or item-item relations ^6^. On the other hand, at least some degree of segregation within the hippocampus has been reported ^10^. Item-item relations may be supported by more anterior regions ^11^, whereas item-space relations may be supported more strongly by right-lateralized posterior hippocampal regions ^12^. Here, we adopt a developmental approach to address the question of whether developmental improvements in these three forms of relational memory rely on structural changes in the hippocampus and, if so, whether they depend on the same or different subregions.

Recent research has highlighted age-related differences in hippocampal structure and function in children and adolescents and evidence of cross-sectional associations between volume and memory ^13–16^. However, longitudinal evidence linking changes in hippocampal structure to memory development is lacking. We shed new light on these issues by capitalizing on a longitudinal design in which we assessed both structural changes in hippocampal head, body, and tail subregions and behavioral changes in an experimental task assessing item-space, item-time and item-item memory.

There are at least two lines of evidence suggesting that this approach may be particularly informative. First, initial cross-sectional findings suggested heterogeneous development of the hippocampus along the anterior-posterior axis with distinct relations with memory^14,16–18^. Second, heterogeneities in age-related differences in memory for spatial, temporal and associative information have been documented in cross-sectional studies against a backdrop of general memory improvement during childhood ^15,19–21^. This body of research indicates that memory for spatial relations may be more robust at a younger age compared to memory for temporal relations ^20–22^ and item-item associative relations ^22^. Overall, these two lines of evidence suggest a co-occurrence of distinct structural changes in the anterior and posterior hippocampus and distinct behavioral changes in relational memory, consistent with a functional segregation in the hippocampus during development. However, an important limitation of these cross-sectional studies is that it was not possible to examine whether developmental changes in hippocampal structures predicted developmental improvements in memory over time within the same individuals.

In the present study, we used a combination of experimental and longitudinal approaches to examine a cohort of 172 children between 7 and 15 years of age who underwent structural magnetic resonance imaging (MRI) and relational memory assessment on up to three measurement occasions (T1, T2, T3) (Fig. 1A; 362 longitudinal scans; 393 longitudinal behavioral assessments). The advantage of a longitudinal approach combining behavior and brain assessment is its potential to reveal how structural changes predict behavioral development, accounting for concurrent associations. Participants encoded triplets of novel visual objects, each appearing one at a time in one of three locations on the screen (Figure 1B, Top). Memory was tested immediately after study with a probe signaling whether participants were required to retrieve item-space, item-time, or item-item associations (Figure 1B, Bottom).

The central hypothesis guiding the present research is that changes in hippocampal structure contribute to developmental improvements in relational memory. Specifically, we predicted that relational memory developed differentially as a function of type of relation, with the ability to remember item-space relations developing earlier than the other relations. We also predicted distinct developmental trajectories of hippocampal volume as a function of subregion, with the hippocampal head decreasing and the hippocampal body increasing in volume at least prior to age 10 ^15^. Finally, we hypothesized that volumetric changes in hippocampal subregions would predict behavioral changes differently as a function of type of relation. For example, changes in more posterior subregions (i.e., tail) were expected to relate to the development of memory for item-space relations ^10^.

**Figure 1.**
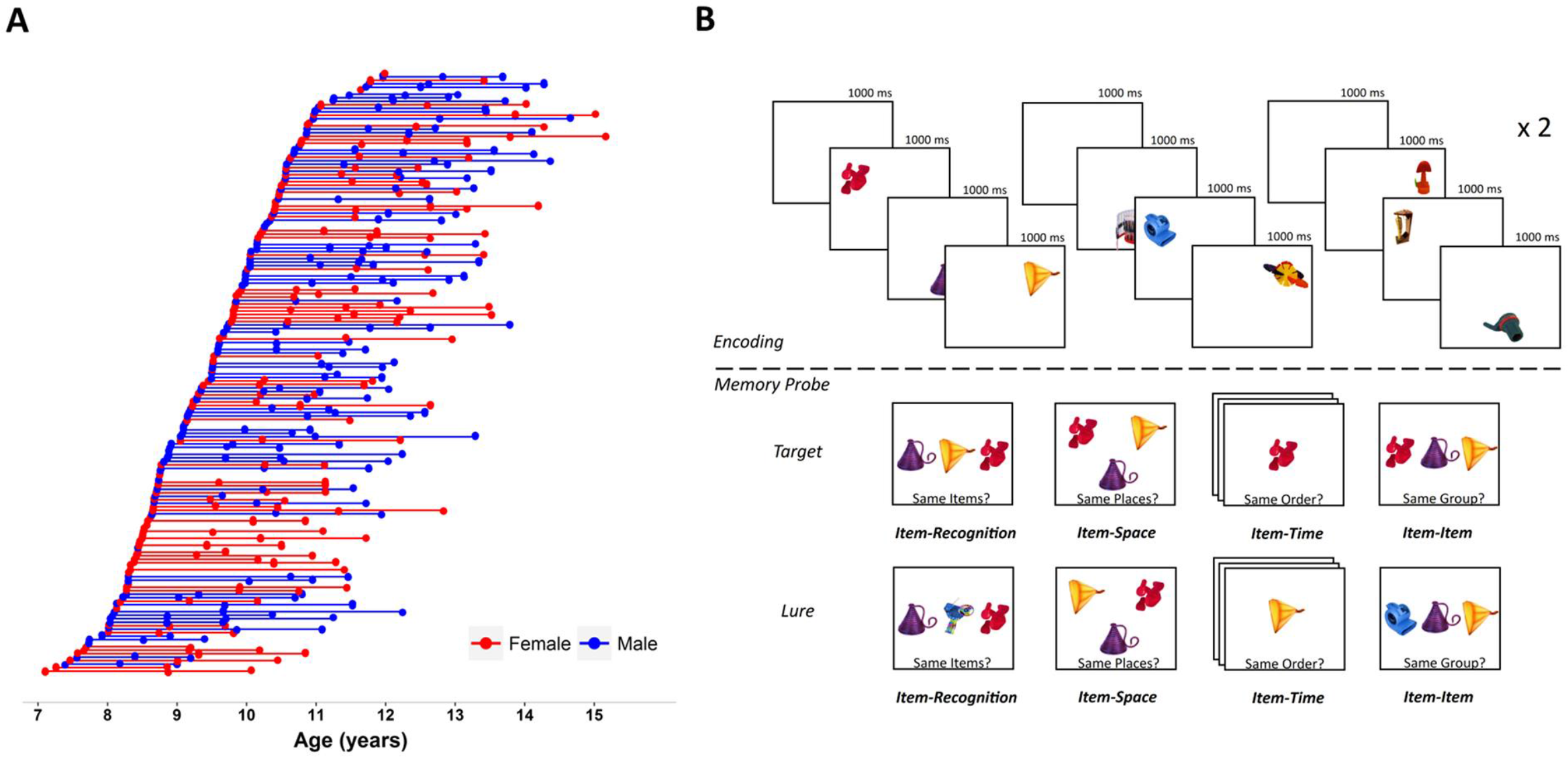
**A.** Longitudinal cohort of 172 children providing MRI structural images and relational memory assessments on up to three occasions (362 longitudinal scans, 393 longitudinal behavioral assessments). **B.** Triplet Binding Task (TBT). Encoding: Item-Recognition, Item-Space, Item-Time, and Item-Item relation conditions shared identical encoding procedures. Memory probe: Target and lure test trials for item-recognition, item-space, item-time, and item-item relation conditions, from left to right, respectively.

To briefly summarize our key and novel findings, we report that memory for item-space relations matured earlier than memory for item-time and item-item relations, and that the hippocampal head declined in volume throughout most of middle childhood, whereas hippocampal body increased in volume until approximately age 10 before declining. Finally, we report that volumetric increases in head and body predicted better item-time and item-time memory, whereas increases in tail volume predicted better item-space memory.

## Results

We conducted longitudinal analyses using mixed effect models ^23^. Memory for each relation was calculated as the difference between hit and false-alarm rates. Total hippocampal volumes were first extracted using the semi-automated procedure described in the Methods section, and were then manually segmented into head, body and tail based on established guidelines ^14^. This segmentation had excellent inter-rater reliability (Head/Body Division: ICC=.98; Body/Tail Division: ICC=.99). Volumes were adjusted for intracranial volume (ICV) using regression methods ^24^. In all models, the effect of age was separated into a time-varying within-subject effect (i.e., change in age since T1) and a time-invariant between-subject effect (i.e., age at T1) (25, 27; see Methods). In brain–behavior models, the effects of head, body, and tail volumes were similarly separated into a time-varying within-subject effect (i.e., changes in volume since T1) and a time-invariant between-subject effect (i.e., volume at T1).

In each longitudinal analysis, model comparisons were conducted to test whether the inclusion of key variables of interest increased model fit over baseline models, beginning with testing for main effects, and then systematically adding higher order interaction effects with these key variables. The full longitudinal models are described in Table 1. The key variables of interest in the behavioral models included the effect of age at T1 and change in age, as well as the two-way interactions between these variables and three-way interactions with type of memory relation. The key variables of interest in the hippocampal models were also age at T1 and change in age, as well as their interaction, and three-way interactions with hippocampal subregion. Finally, in the brain–behavior models, the key variables of interest were volume of head, body, and tail at T1 and changes in these volumes since T1, as well as their interactions with age at T1 and change in age.

**Table 1.**
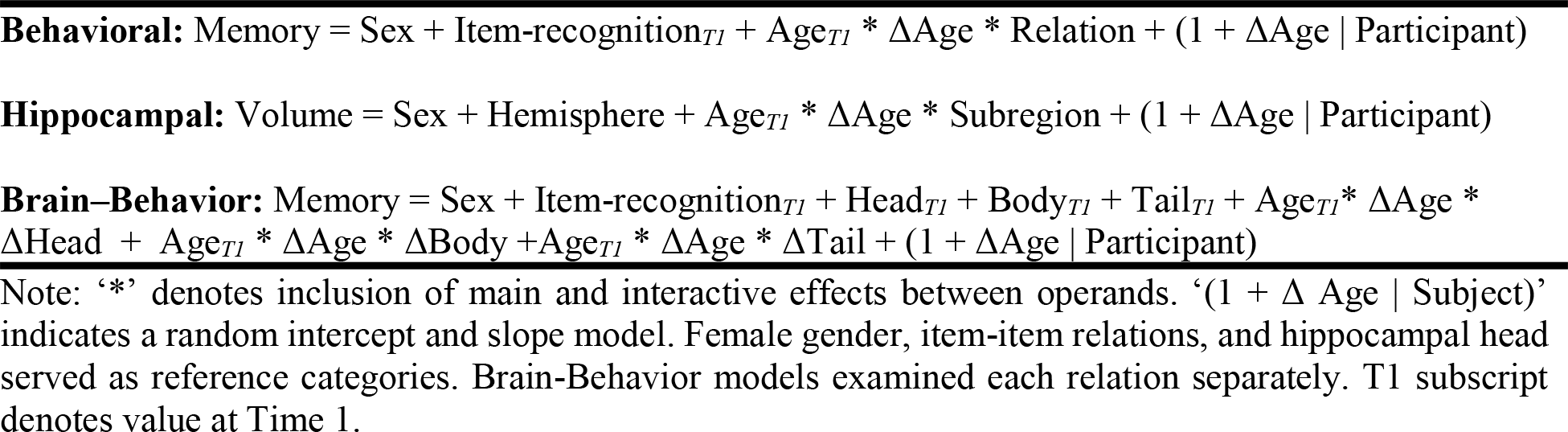
Fixed and Random Effect Models

### Distinct Developmental Trajectories of Relational Memory

We first conducted the longitudinal analysis of relational memory (See Table 1). Overall, relational memory was greater in children who were older at T1 (χ2 = 17.8, *df* = 1, *p* <.0001; β=.18, *b* = .04, *t*(170) = 4.4, *p* <.0001), capturing cross-sectional differences, and it increased more as more time passed, as indicated by a positive association with change in age (χ2 = 25.5 *df* = 1, *p* <.0001; β=.17, b=.04, *t*(121)=5.19, *p* <.0001). Improvements in relational memory over time were greater for children who were younger at T1 (age at T1 × change in age in years interaction; χ2 = 7.90, *df* = 1, *p* = .005; β=.18, b=−.02, *t*(140)=−2.88, *p* = .004). We also found a significant effect of type of relation (χ2 = 368.5, *df* = 2, *p* <.0001), such that the highest performance was observed for item-space memory (M=.45; SE= .01), which was greater than item-time (M=.36, SE=.01; *t* (864) = 7.1, *p* <.0001). Item-time was, in turn, greater than item-item memory (M=.17, SE .01; *t* (864) = 10.03, *p* < .0001). Consistent with our primary hypothesis, the magnitude of memory improvement over time depended on the type of relation, as indicated by a significant interaction between change in age and type of relation (χ2 = 6.21 *df* = 2, *p* = .04) (Figure 2). See Table 2 for parameter estimates for each type of relation separately, and Table S1 for parameter estimates testing the interaction with type of relation. The positive association between change in age and change in memory was stronger for item-time and item-item than for item-space (item-space: β=.09, *b* = .02, *t* (374) = 2.17, *p* = .03; item-time relative to item-space, β=.08, *b* = .03, *t* (867) = 2.18, *p* = .03; item-item relative to item-space, β=.08, *b* =.03, *t* (867) = 2.11, *p* = .04). Associations between change in age and performance did not differ between item-time and item-item relations (*p* =.94). Model parameters predicted that item-space memory plateaued around 10.4 years, item-time memory around 12.2 years of age, and item-item around 12.5 years. Thus, consistent with prior work, item-space memory matured earlier than both item-item and item-time relations.

**Figure 2.**
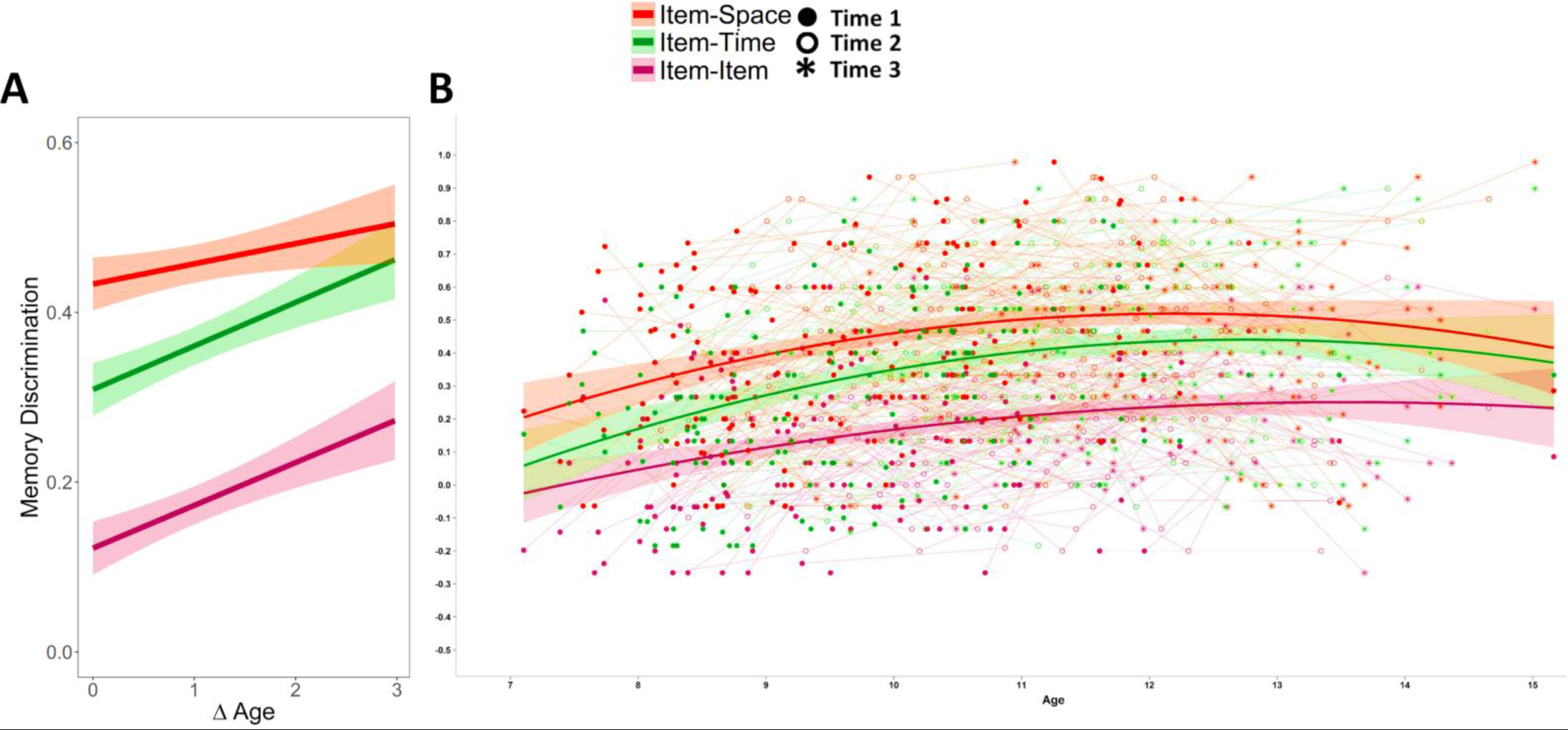
Developmental changes in memory for item-space, item-time, and item-item relations. Error bands represent 95% confidence intervals. **A.** Depicting the three-way interaction between memory relation, within-subject changes in age since Time 1 (ΔAge), and cross-sectional differences in the starting age at Time 1 (here at 8- and 11-years of age). **B.** A descriptive spaghetti plot of item-space, item-time, and item-item memory performance by years in age, with quadratic lines fitted. Note that the use of age conflates between-person cross-sectional differences with within-person changes, and thus these fit lines do not reflect true longitudinal change.

**Table 2.** Parameter Estimates for Item-Time, Item-Item and Item-Space Models

| Effect | Beta | b | SE | t | p |
| --- | --- | --- | --- | --- | --- |
| **Item-Time** |  |  |  |  |  |
| (Intercept) | – | .323 | .023 | 14.3 | <.001 |
| Item-Recognition | .310 | .353 | .066 | 5.39 | <.001 |
| Male | -.048 | -.025 | .029 | -.861 | .390 |
| Start-Age | .213 | .044 | .013 | 3.29 | .001 |
| ΔAge | .212 | .051 | .011 | 4.61 | <.001 |
| Start-Age x ΔAge | -.125 | -.019 | .009 | -2.05 | .043 |
| **Item-Item** |  |  |  |  |  |
| (Intercept) | – | .133 | .019 | 6.93 | <.001 |
| Item-Recognition | .162 | .151 | .053 | 2.87 | .005 |
| Male | -.033 | -.014 | .023 | -.605 | .546 |
| Start-Age | .204 | .035 | .012 | 2.95 | .004 |
| ΔAge | .244 | .048 | .009 | 5.27 | <.001 |
| Start-Age x ΔAge | -.128 | -.016 | .008 | -2.07 | .041 |
| **Item-Space** |  |  |  |  |  |
| (Intercept) | – | .457 | .023 | 20.2 | <.001 |
| Item-Recognition | .328 | .357 | .065 | 5.49 | <.001 |
| Male | -.076 | -.038 | .029 | -1.31 | .191 |
| Start-Age | .180 | .036 | .014 | 2.66 | .009 |
| ΔAge | .083 | .019 | .011 | 1.73 | .086 |
| Start-Age x ΔAge | -.139 | -.020 | .009 | -2.18 | .031 |

Notes: Model Fits: Item-Time: $\chi^2 = 68.7$ , $df = 5$ , $p < 1.85\text{e-}13$ ; Item-Space: $\chi^2 = 48.2$ , $df=5$ , $p = 3.3\text{e-}9$ ; Item-Item: $\chi^2 = 48.0$ , $df=5$ , $p = 3.6\text{e-}9$ ; Interactions with sex were not significant ( $\chi^2 \leq 4.6$ , $dfs=3$ , $ps \geq .20$ ). Note: ΔAge is defined at time in years since Time 1. Item-recognition and Start-Age are centered at the mean at Time 1. Left hemisphere and female are reference categories.

### Distinct Developmental Trajectories of Hippocampal Subregions

We assessed developmental changes in hippocampal head, body, and tail (See Table 1). We found a significant interaction between change in age and hippocampal subregion (χ2 = 8.83 *df* = 2, *p* = .012), which was further moderated by age at T1 (χ2 = 9.80, *df* = 3, *p* = .020). As predicted, we found distinct within-subject trajectories for the three subregions (Figure 3). See Table S2 for parameter estimates of this full model. For completion, we also estimated longitudinal models using total hippocampal volume, the results of which are reported in Table S3. Given the differences in volumetric change as a function of subregion, we examined the trajectory of each subregion separately.

**Figure 3.**
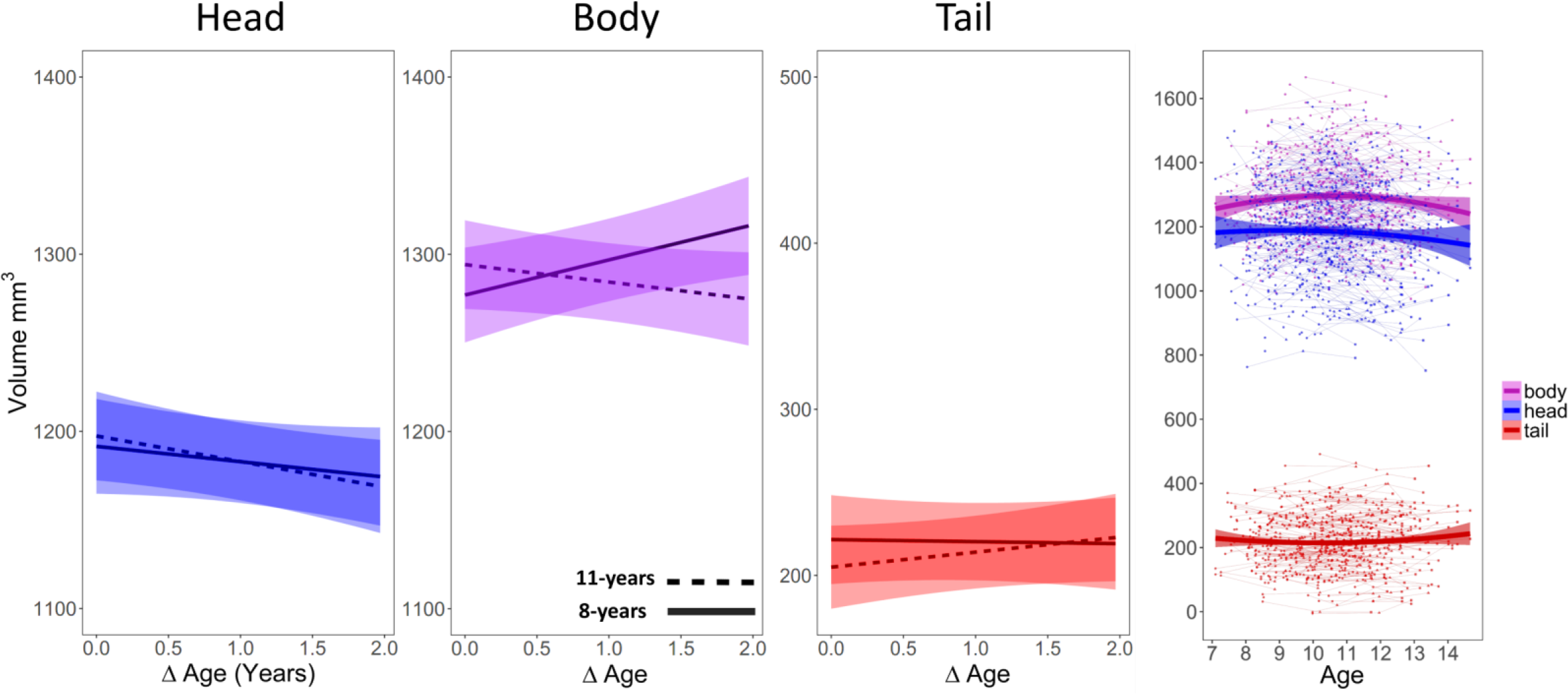
Developmental changes in head, body, and tail ICV-corrected volume. Error bands represent 95% confidence intervals. **A.** Depicting the three-way interaction between hippocampal sub-region, within-subject change in age since Time 1 (ΔAge), and cross-sectional differences in the starting age at Time 1 at 8- and 11-years of age. **B.** Spaghetti plots of head, body, and tail ICV-corrected volume over time with quadratic lines fitted.

**Table 3.** Parameter Estimates for Models of Hippocampal Head, Body, and Tail Change

| Sub-Region | Effect | Beta | b | SE | t | p |
| --- | --- | --- | --- | --- | --- | --- |
| Head | (Intercept) | – | 1128 | 16.4 | 68.8 | <.001 |
|  | Male | .030 | 11.7 | 22.3 | .525 | .600 |
|  | Hemisphere [Right] | .313 | 106 | 5.23 | 20.2 | <.001 |
|  | Start-Age (Mean-Centered) | .011 | 3.02 | 10.2 | .298 | .770 |
| | ΔAge | -.060 | -7.07 | 2.70 | -2.62 | .009 |
| | Start-Age x ΔAge | -.056 | -5.51 | 2.56 | -2.16 | .033 |
| Body | (Intercept) | – | 1314 | 13.4 | 98.1 | <.001 |
|  | Male | -.133 | -26.1 | 18.1 | -1.44 | .150 |
|  | Hemisphere [Right] | -.104 | -33.5 | 4.93 | -6.80 | <.001 |
|  | Start-Age (Mean-Centered) | .015 | 1.40 | 8.35 | .167 | .873 |
| | ΔAge | .012 | 1.68 | 2.53 | .661 | .514 |
| | Start-Age x ΔAge | -.061 | -4.86 | 2.39 | -2.03 | .042 |
| Tail | (Intercept) | – | 208 | 9.38 | 22.1 | <.001 |
|  | Male | .067 | 11.5 | 12.6 | .912 | .363 |
|  | Hemisphere [Right] | .024 | 4.10 | 2.92 | 1.40 | .164 |
|  | Start-Age (Mean-Centered) | -.042 | -3.30 | 5.86 | -.564 | .572 |
| | ΔAge | .022 | 1.76 | 1.50 | 1.17 | .240 |
| | Start-Age x ΔAge | .010 | .538 | 1.42 | .377 | .712 |

Model Fits: Hippocampal Head: $\chi^2 = 312$ , $df = 5$ , $p < 2.2e-1$ ; Hippocampal Body: $\chi^2 = 51.4$ , $df=5$ , $p = 7.2e-10$ ; Hippocampal Tail: $\chi^2 = 4.44$ , $df= 5$ , $p = .49$ . Note: ΔAge is defined as time in years since Time 1; Left hemisphere and female are reference categories; Volumes are in cubic mm.

#### Hippocampal Head

As predicted, hippocampal head volumes declined over time, as indicated by the negative effect of change in age (χ2 = 5.63, *df* = 1, *p* = .02; *b* = −7.07, *t* (449) = −2.62, *p* = 9.2e-3). This effect was moderated by age at T1 (χ2 = 4.65, *df* = 1, *p* = .03; β=−.06, *b* =−5.51, *t* (457) = −2.16, *p* = .03), such that greater volumetric declines were observed in children the older you were at T1. Associations with change in age did not significantly differ between hemispheres (χ2 = .60, *df* = 1, *p* = .44) or sex (χ2 = 2.58, *df* = 1, *p* = .11) (Table 3). A descriptive examination of the partial derivatives of model parameters suggests that peak volume of hippocampal head occurred at 8.17 years of age before declining during late childhood.

#### Hippocampal Body

As predicted, hippocampal body exhibited a non-linear trajectory. Change in age significantly interacted with age at T1 (χ2 = 4.10, *df* = 1, *p* = .04; β=−.06, *b* = −4.86, *t* (496) = −2.03, *p* = .04): The volume of the hippocampal body increased over time for younger children, but it declined for older children. Association with changes in age did not significantly differ by hemisphere (χ2 = .60, *df* = 1, *p* = .44) or sex (χ2 = 3.4e-3, *df* = 1, *p* = .95) (Table 3). A descriptive examination of the partial derivatives of model parameters suggests that peak volume of hippocampal body occurred at 9.79 years before declining in late childhood.

#### Hippocampal Tail

No significant developmental changes were observed for either left or right tail (Table 3).

### Linking Hippocampal and Relational Memory Development

We examined whether and how volumetric changes along the anterior-posterior axis predicted the development of each type of memory relation (See Table 1). All models included volume at T1, changes in volume since T1, age at T1, and changes in age since T1, as well as their interactions. Volume and volume changes were in cubic millimeters for unstandardized betas. The primary longitudinal effects of interest were the two- and three-way interactions between age at T1, change in age, and change in volume. These interactions allow us to link developmental changes in volume to behavioral development, with the additional consideration that longitudinal relations may depend on the age at the start of the study. We started by examining item-time and item-item memory, because they showed the most robust behavioral change, and ended with item-space memory, which we established develops relatively earlier (see Methods for detailed description of the models). For these, left and right hippocampal volumes were summed because no hemispheric differences were observed.

#### Item-Time

Consistent with predictions, changes in hippocampal head, body, and tail predicted item-time memory. Specifically, we observed a significant three-way interaction between change in hippocampal subregion volumes, age at T1 and change in age (χ2 = 12.1, *df* =3, *p* = .007) (See Table 4). Increase in head and body volumes, but not tail, significantly predicted greater memory performance after longer delays (e.g., a 3-year change is depicted in Figure 4A), but not shorter delays (e.g., a 1-year change in age is depicted in Figure S1A), indicating that several years were necessary for these brain-behavior relations to manifest. Furthermore, this result depended on age at T1. When the model was evaluated for children who were older at T1 (e.g., 11 years, as depicted in Figure 4A), volumetric increases in head and body volume predicted better item-time memory (Body: β=.47, b=.001, *SE* = 4.9e-4, *t*=2.59, *p*=.01; Head: β=.35, *b*=.001, *SE* = 5.1e-4, *t*=1.87, *p*=.06), but was not significant for children who were younger at T1 (e.g., 8 years, as depicted in Figure 4A), despite the appearance of a negative relation (*p*s ≥.17). Change in the tail was not associated with item-time performance (*p*s ≥ .18). Thus, although the normative pattern of volumetric change in this sample was a linear decrease in the head, and a curvilinear in the body volume over time, protracted increases in head and body volume in older children predicted better item-time memory. Parameter estimates for models separating left and right hippocampal structures are also included in Table S4.

**Table 4.** Hippocampal Volume Predicting the Development of Item-Time Memory.

| Effect | Left and Right Hippocampal Sum |  |  |  |  |
| --- | --- | --- | --- | --- | --- |
|  | Beta | b | SE | t | p |
| (Intercept) | - | 3.2e-1 | 2.5e-2 | 13 | <.0001 |
| Item-Recognition | 0.28 | 3.2e-1 | 7.4e-2 | 4.4 | <.0001 |
| Sex [Male] | -0.053 | -2.8e-2 | 3.2e-2 | -0.86 | 0.39 |
| Start-Volume Head | -0.049 | -4.5e-5 | 5.6e-5 | -0.79 | 0.43 |
| Start-Volume Body | -0.065 | -7.0e-5 | 6.6e-5 | -1.1 | 0.29 |
| Start-Volume Tail | 0.062 | 9.8e-5 | 9.9e-5 | 0.99 | 0.32 |
| Start-Age | 0.25 | 5.8e-2 | 1.7e-2 | 3.4 | 0.001 |
| ΔAge | 0.26 | 6.2e-2 | 1.3e-2 | 4.7 | <.0001 |
| ΔHead | -0.063 | -1.9e-4 | 4.1e-4 | -0.47 | 0.64 |
| ΔBody | -0.056 | -1.6e-4 | 3.8e-4 | -0.43 | 0.67 |
| ΔTail | -0.2 | -1.4e-3 | 9.5e-4 | -1.5 | 0.14 |
| Start-Age x ΔAge | -0.13 | -2.1e-2 | 1.2e-2 | -1.7 | 0.095 |
| Start-Age x ΔHead | -0.26 | -6.4e-4 | 3.2e-4 | -2 | 0.048 |
| Start-Age x ΔBody | -0.22 | -6.3e-4 | 3.7e-4 | -1.7 | 0.096 |
| Start-Age x ΔTail | -0.037 | -2.6e-4 | 1.0e-3 | -0.25 | 0.8 |
| ΔAge x ΔHead | 0.072 | 1.1e-4 | 2.2e-4 | 0.5 | 0.62 |
| ΔAge x ΔBody | 0.14 | 2.0e-4 | 1.9e-4 | 1.1 | 0.29 |
| ΔAge x ΔTail | 0.11 | 4.0e-4 | 5.0e-4 | 0.8 | 0.42 |
| Start-Age x ΔAge x ΔHead | 0.33 | 4.1e-4 | 1.9e-4 | 2.2 | 0.027 |
| Start-Age x ΔAge x ΔBody | 0.29 | 4.2e-4 | 1.9e-4 | 2.2 | 0.032 |
| Start-Age x ΔAge x ΔTail | 0.12 | 4.2e-4 | 5.5e-4 | 0.77 | 0.44 |

Note: Female is reference sex. For unstandardized betas, volume is in cubic millimeters and age is in years.

**Figure 4.**
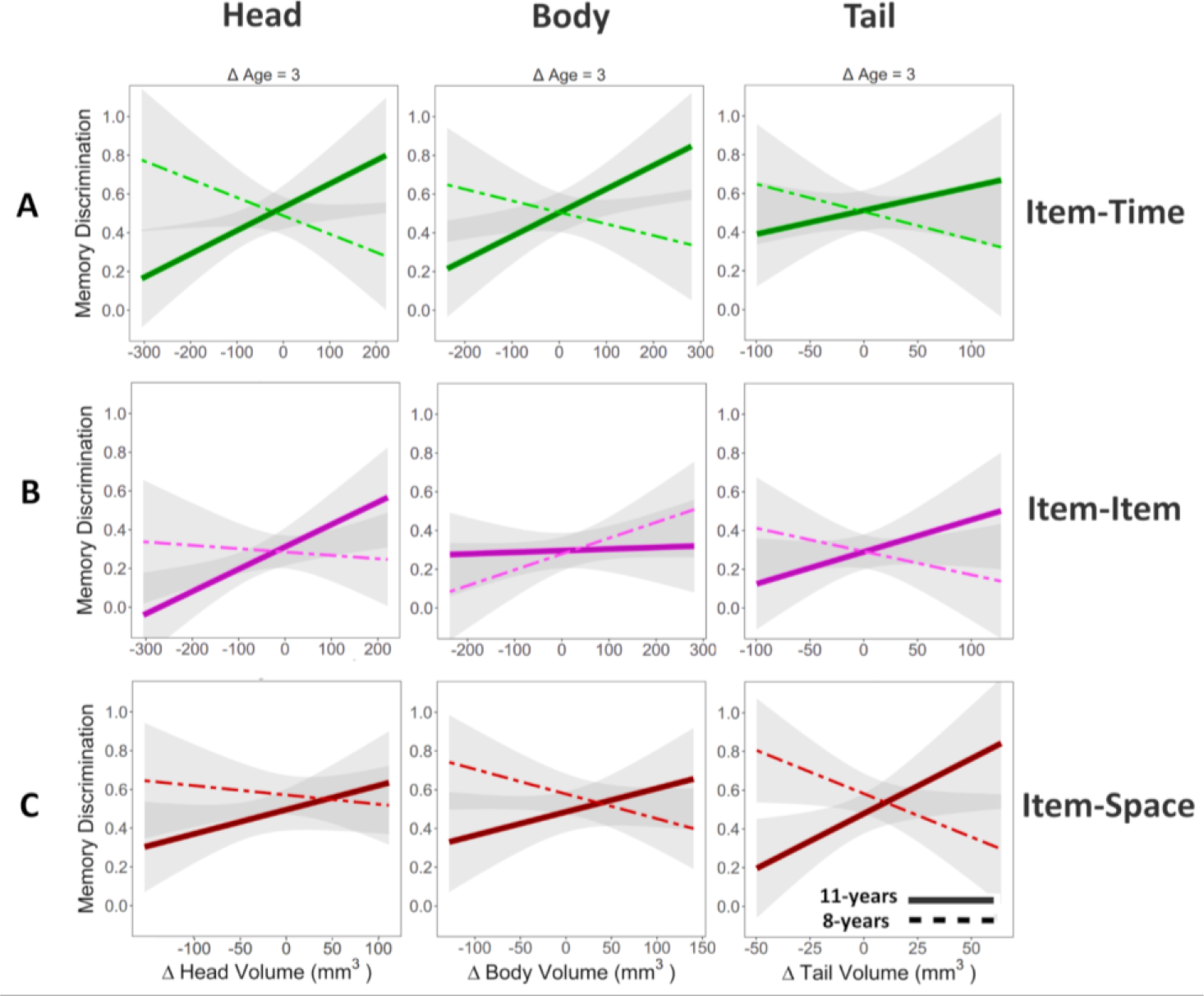
Depicting interaction between change in ICV-corrected volume and cross-sectional differences in the starting age at Time 1 at 8- and 11-years of age evaluated at change in age since Time 1 equaling three years (ΔAge = 3). See Supplemental Figure 1 for depiction of interaction after one year since Time 1; relations between volume changes and memory were stronger at longer delays. Error bands represent 95% confidence intervals. **A.** Item-Time. **B.** Item-Item. **C.** Item-Space.

#### Item-Item

Consistent with our prediction, changes in hippocampal structure predicted item-item memory. Specifically, we found a significant interaction between volumetric changes in head, body, and tail (as a block) and age at T1 (χ2 = 8.82, *df* =3, *p* =.03), but this interaction was not significantly moderated by changes in age (χ2 = 3.2, *df* = 3, *p* = .37) (See Table 5). Examining the volumetric change and age at T1 interaction, we found that among children who were young at T1 (i.e., 8 years), increases in body volume predicted greater item-item memory (β=.27, *b*=.0007, *SE* = 2.5e-4, *t*=2.93, *p*=.004). In contrast, among children who were older at T1 (i.e., 11 years), increases in head volume predicted better behavioral performance (β=.24, *b*=.0006, *SE* = 2.3e-4, *t*=2.38, *p*=.02) (See Figure 4B and Figure S1B). Parameter estimates for models separating left and right hippocampal structures are also included in Table S5.

**Table 5.**
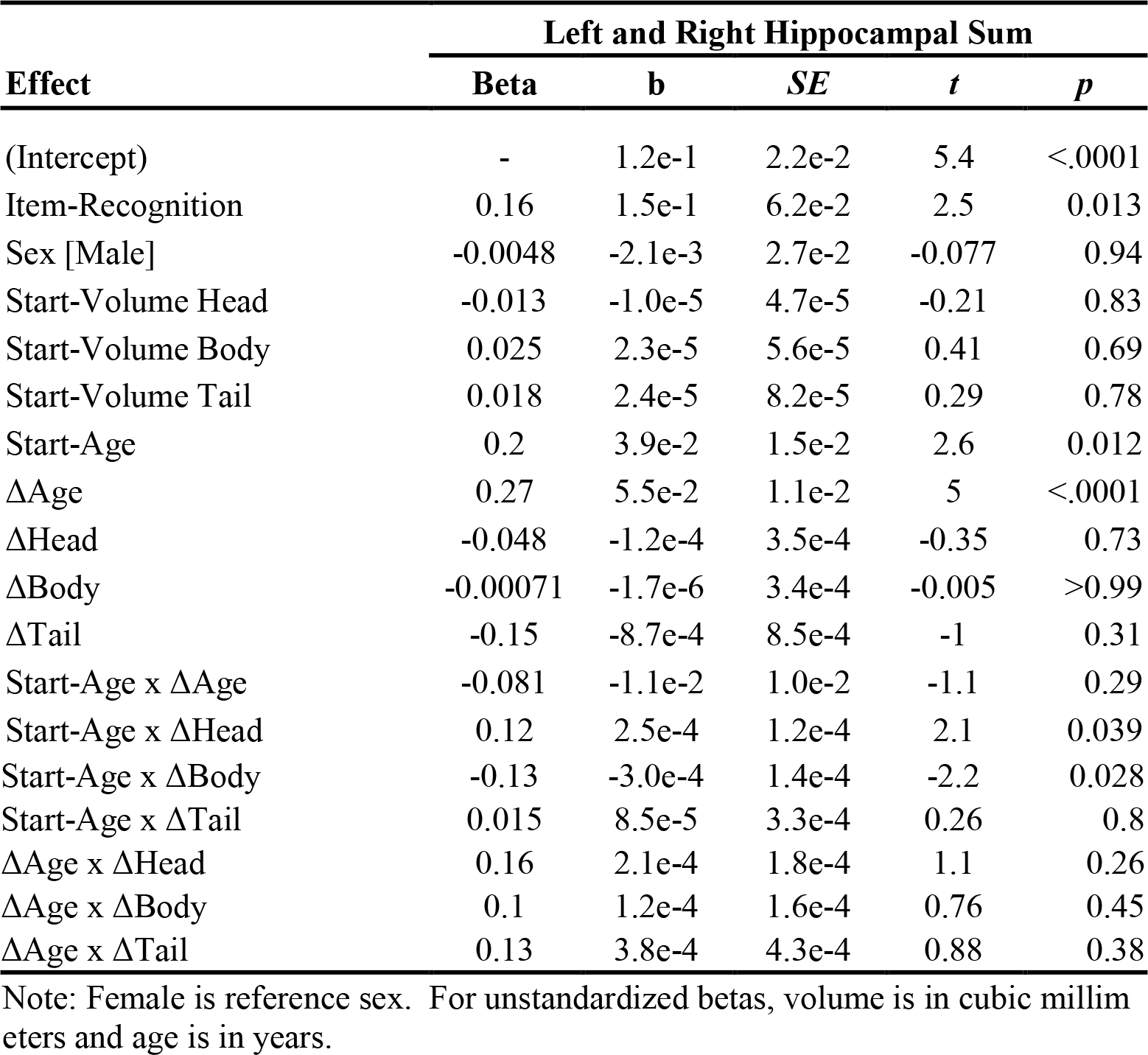
Hippocampal Volume Predicting the Development of Item-Item Memory.

Overall, volumetric changes in hippocampal body appeared to differentially predict item-time and item-item memory. Consistent with this, we found that the age at T1 by change in body volume interaction was significantly different for item-time and item-item memory (χ2 = 8.92, *df* = 1, *p* = .003). In younger children, the association between change in body and memory was more positive for item-item than item-time (β=.32, *b*=001, *SE* = 5.2e-4, *t*=2.50, *p*=.014), but in older children, there was a trend for a more negative relation for item-item than item-time memory (β=−.28, *b*=−.001, *SE* = 5.8e-4, *t*=−1.93, *p*=.055). Overall results are consistent with the protracted behavioral trajectory of item-item memory and suggest a transition from body to head in supporting developmental improvements in item-item memory.

#### Item-Space

No significant relations between changes in hippocampal structure and item-space memory were found when we used volume changes summed across hemispheres (χ2s ≤ 4.04, *df*s = 3, *p*s ≥ .26) (See Table S6), nor did using overall hippocampal volume perform better than using subregions (χ2 = 3.84, *df* = 8, *p* = .87).

Given the suggestion from the literature that associations between change in head, body, and tail volumes and spatial memory could be right-lateralized, we also tested our model in the right hippocampus. This analysis revealed a significant three-way interaction between changes in right hippocampus, changes in age, and starting age at T1 (χ2 = 10.6, df = 3, *p* = .01) (See Table 6). Volumetric changes significantly more positively predicted memory performance with longer delay (e.g. 3 years; Figure 4C), but not significantly with shorter delays (e.g., 1 year; *p*s > .098; Figure S1C). In other words, in younger children at T1, there was a trend for reduction of tail volume over time predicting better item-space memory (β=−.32, b= −.004, *SE* = .002, *t*=−1.86, *p*=.07), but in older children at T1, volumetric increases in the tail predicted better item-space memory (β=.528, b=.006, *SE* = .003, *t*=2.16, *p*=.03). However, neither the body (*p*s ≥ .11) nor the head (*p*s ≥ .21) were significantly associated to item-space memory at those starting ages. Thus, although the hippocampal tail did not seem to show an average pattern of volumetric change based on previous analyses, the present results suggest that individual differences in tail development predict item-space memory performance.

**Table 6.**
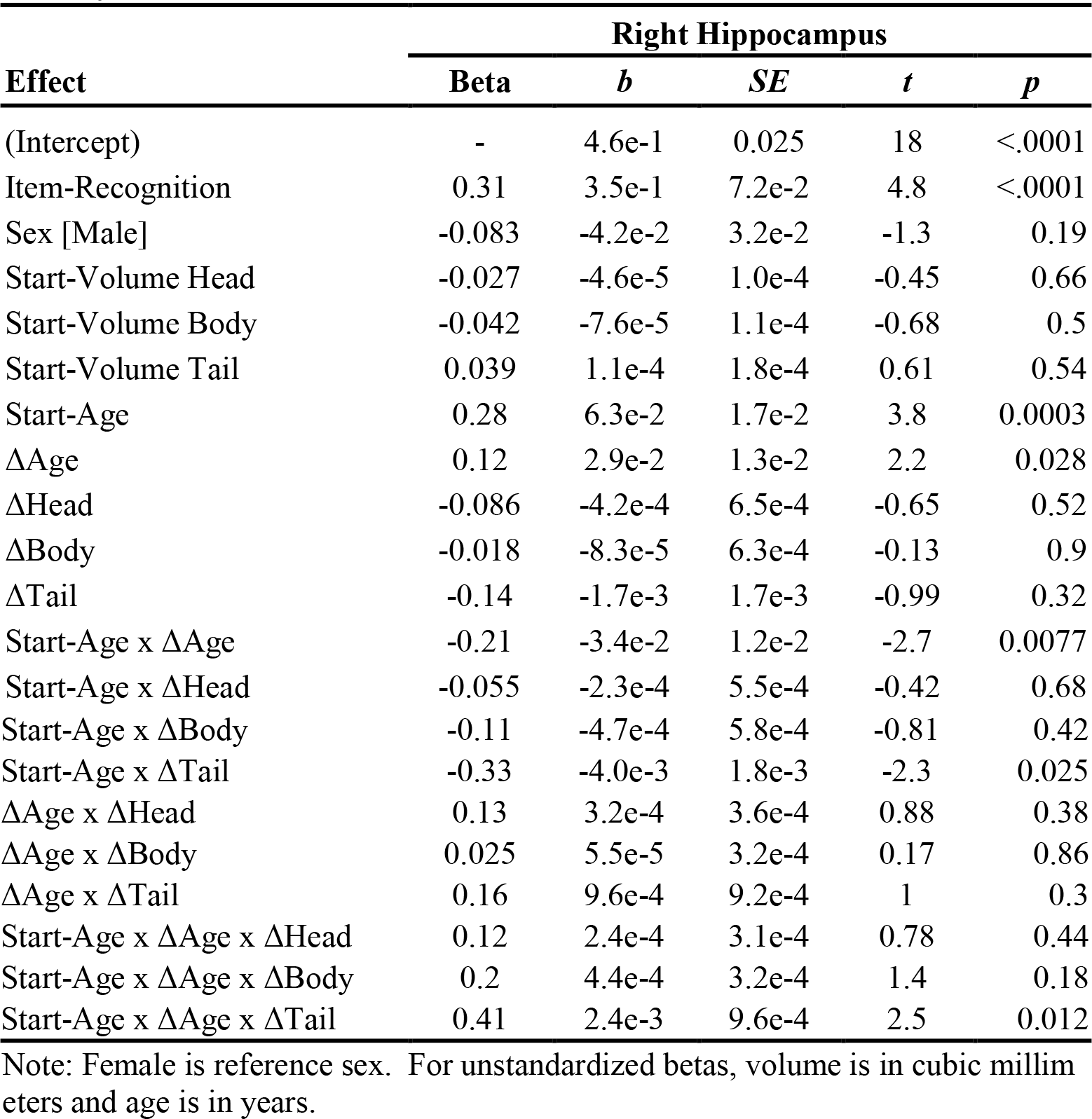
Hippocampal Volume Predicting the Development of Item-Space Memory.

| Effect | Right Hippocampus |  |  |  |  |
| --- | --- | --- | --- | --- | --- |
|  | Beta | *b* | *SE* | *t* | *p* |
| (Intercept) | - | 4.6e-1 | 0.025 | 18 | <.0001 |
| Item-Recognition | 0.31 | 3.5e-1 | 7.2e-2 | 4.8 | <.0001 |
| Sex [Male] | -0.083 | -4.2e-2 | 3.2e-2 | -1.3 | 0.19 |
| Start-Volume Head | -0.027 | -4.6e-5 | 1.0e-4 | -0.45 | 0.66 |
| Start-Volume Body | -0.042 | -7.6e-5 | 1.1e-4 | -0.68 | 0.5 |
| Start-Volume Tail | 0.039 | 1.1e-4 | 1.8e-4 | 0.61 | 0.54 |
| Start-Age | 0.28 | 6.3e-2 | 1.7e-2 | 3.8 | 0.0003 |
| ΔAge | 0.12 | 2.9e-2 | 1.3e-2 | 2.2 | 0.028 |
| ΔHead | -0.086 | -4.2e-4 | 6.5e-4 | -0.65 | 0.52 |
| ΔBody | -0.018 | -8.3e-5 | 6.3e-4 | -0.13 | 0.9 |
| ΔTail | -0.14 | -1.7e-3 | 1.7e-3 | -0.99 | 0.32 |
| Start-Age x ΔAge | -0.21 | -3.4e-2 | 1.2e-2 | -2.7 | 0.0077 |
| Start-Age x ΔHead | -0.055 | -2.3e-4 | 5.5e-4 | -0.42 | 0.68 |
| Start-Age x ΔBody | -0.11 | -4.7e-4 | 5.8e-4 | -0.81 | 0.42 |
| Start-Age x ΔTail | -0.33 | -4.0e-3 | 1.8e-3 | -2.3 | 0.025 |
| ΔAge x ΔHead | 0.13 | 3.2e-4 | 3.6e-4 | 0.88 | 0.38 |
| ΔAge x ΔBody | 0.025 | 5.5e-5 | 3.2e-4 | 0.17 | 0.86 |
| ΔAge x ΔTail | 0.16 | 9.6e-4 | 9.2e-4 | 1 | 0.3 |
| Start-Age x ΔAge x ΔHead | 0.12 | 2.4e-4 | 3.1e-4 | 0.78 | 0.44 |
| Start-Age x ΔAge x ΔBody | 0.2 | 4.4e-4 | 3.2e-4 | 1.4 | 0.18 |
| Start-Age x ΔAge x ΔTail | 0.41 | 2.4e-3 | 9.6e-4 | 2.5 | 0.012 |

Note: Female is reference sex. For unstandardized betas, volume is in cubic millimeters and age is in years.

## Discussion

The ability to remember associations between events and their spatio-temporal context depends on hippocampal mechanisms, which bind contextual features into integrated event representations ^1^. Here, we asked whether volumetric changes in hippocampal volume predict longitudinal improvements in relational memory, and whether those developmental associations differed depending on hippocampal subregion or type of memory relation.

This is the first report showing that longitudinal improvements in relational memory differed as a function of the type of memory relation, such that item-space memory developed more rapidly than item-time and item-item memory. In the largest longitudinal study of hippocampal subregions to date, this research showed that hippocampal head, body, and tail follow different developmental trajectories from childhood into adolescence. Linking structural and behavioral changes, we report for the first time that volumetric changes in hippocampal head, body, and tail differentially predicted longitudinal improvement in item-space, item-time, and item-item.

### Developmental Change in Relational Memory Depends on the Nature of the Relation

In our initial cross-sectional analysis ^22^, item-space memory reached adults’ levels of performance before item-time memory, which in turn preceded item-item memory. In the present research, we examined within-person change while accounting for cross-sectional differences and showed that item-space memory improves until around 10½, whereas item-time and item-item memory followed prolonged trajectories with improvements about 12 and 12½ years of age respectively. This finding is additionally consistent with prior cross-sectional evidence that spatial memory develops earlier than temporal memory ^20–22^. Although we cannot rule out the possibility that aspects of our tasks might differ across conditions for reasons other than the type of relation manipulated, we argue that the use of novel stimuli and arbitrary associations is an effective way to assess relational memory. The more rapid development of item-space memory compared to the other relations suggests that relational memory processes are not fully unitary.

Although item-time memory was generally better than item-item, their developmental trajectories were similar. This may have been due to the dependence of these tasks on shared hippocampal operations. For example, performance on both item-time and item-item memory may have benefitted from some form of temporal processing—the former from processing the precise temporal order of the images and the latter from processing which groups of items were presented together in the same temporal context ^7^. On the other hand, there may also be differences in how the hippocampus supports item-time and item-item memory despite the apparent similarity in behavioral trajectory, which may help to explain why item-item is a more challenging task ^26,27^. Disentangling these two possibilities was made possible by the longitudinal design combining assessments of both brain and behavior and was addressed in the brain–behavior analyses. Overall, these behavioral findings provide the first longitudinal evidence of protracted and distinct developmental trajectories of different aspects of relational memory. The examination of these relations within participants and within the same task form, which constrain response demands, offers strong support for a functional distinction in relational memory.

### Developmental Change in Hippocampal Volumes Varies Along the Anterior-Posterior Axis

We provided new longitudinal evidence indicating that hippocampal head, body, and tail develop differentially from middle childhood into adolescence. Consistent with the findings of the seminal longitudinal study of 31 individuals that first examined morphometric development along the anterior–posterior axis ^28^, hippocampal head declined in volume from middle childhood to adolescence, while hippocampal body increased in volume until about 10 years of age and declined thereafter. Hippocampal tail volume was stable throughout middle childhood and adolescence, suggesting that its development occurred earliest, consistent with previous reports ^14,16,28^.

Curvilinear trajectories in hippocampal development are frequently observed ^15,18^. Although not yet definitively linked, volumetric increases may reflect ongoing synaptogenesis and dendritic elaboration, while volumetric declines may reflect synaptic pruning ^29^. It is not known why the body, unlike the head and the tail, continues to increase in volume into late childhood (i.e. 9 to 10 years of age). However, the body has been postulated to act as a bridge or integrator of anterior and posterior mechanisms ^30^. We can speculate that continued dendritic elaboration in the body, compared to head and tail, may be important for the body to complete the required connections with head and tail. Whatever the reason, the diverging developmental trajectories of head, body, and tail reported here provide a demonstration that the hippocampus is not a uniform structure and joins the growing body of evidence suggesting functional differences along the anterior–posterior hippocampal axis ^10^.

### Changes in Hippocampal Volume Predict Developmental Improvements in Relational Memory

We found evidence that increases in hippocampal volumes over time predicted longitudinal improvements in relational memory. We note that these positive relations with behavior are observed even in the context of normative volumetric decreases (e.g., hippocampal head). Previous cross-sectional studies have reported negative associations between hippocampal head volume and behavior ^14,17^, suggesting the hypothesis that decreases of hippocampal head over time may promote behavioral improvements. Instead, even though we confirmed normative volumetric declines in this region during development, greater memory performance was observed among those with a relative *increase* in volume. These findings may shed light on underlying mechanisms. One possibility is that these positive associations may depend on ongoing synaptogenesis and dendritic elaboration within hippocampal circuitry ^31^ and these processes may be particularly important for behavior, even when other mechanisms of structural change, such as pruning, may result in a net loss of volume. Our findings overall support a nascent body of cross-sectional research obtained over the last decade linking the hippocampus to age differences in memory ^13,14^. These findings dispel a long-held, but not adequately tested assumption, that the hippocampus and the associative processes it supports, do not contribute to developmental improvements in memory after early childhood ^19^.

We also assessed, for the first time, whether the longitudinal association between hippocampal structure and memory differed as a function of subregion and type of memory relation. These analyses revealed distinct associations, suggesting that processes supporting memory for item-space, item-time, and item-item relations are not uniform across the anterior-posterior axis of the structure. Bilateral increases in the volume of hippocampal head and body predicted larger improvement in item-time memory in older children. In contrast, increases in body volumes predicted item-item memory in younger children and increases in head volume predicted better item-item memory in older children, suggesting a developmental transition from body to head for this type of relation. Finally, the relation between volumetric changes and the development of item-space memory was right lateralized and restricted to the tail, increases in right hippocampal tail over time predicted greater item-space memory, particularly in older children.

Overall, these data suggest that protracted increase in sub-regional volumes are associated with behavioral improvement. It is somewhat surprising that we did not detect reliable relations between hippocampal growth and memory in younger children for item-time and item-space memory. It is possible that memory improvements in younger compared to older children reflect not only change in relational memory, but also increased consistency in children’s engagement with the memory task, potentially obscuring relations between memory and volumetric change. However, contrary to this possibility, we found an association between increases in hippocampal body in younger children and item-item memory, the most difficult of the three relational tasks and, potentially, the most likely to produce less consistent data. Nevertheless, we cannot exclude that our *change in age* parameter captured more variance than our *change in volume parameter* because of additional processing demands in young children. Change in age was included to model time and account for any source of development due to extra hippocampal processes, but shared variance with measures of hippocampal development cannot be excluded.

Our results are consistent with prior evidence that the hippocampus supports memory for item-space, item-time, and item-item relations ^6,8^, but also indicate heterogeneity in how each subregion contributes to these memory relations. Memory for temporal order reliably recruits the hippocampus in functional neuroimaging studies ^3^; however, while we only observed relations with item-time memory for the hippocampal head and body, associations with hippocampal tail have also been reported ^32^, suggesting that temporal memory may not be strongly localized to any anterior-posterior subregion. Memory for associations between items has been preferentially associated with hippocampal head and body ^4,11^, and our results are consistent with these findings. It is notable that item-time and item-item memory trajectories were similar behaviorally. Yet, their trajectories were support by different hippocampal subregions underscoring the advantage of a longitudinal design. Finally, spatial memory is frequently associated with posterior hippocampus (i.e. tail and body) ^12^. We found evidence consistent with this suggestion restricted to the right tail.

Many open questions remain about the processes that might underlie these different longitudinal structure-behavior relations. One possibility is that hippocampal head, body, and tail differ in terms of cell types and genetic expression ^33^, synaptic plasticity ^34^, and relative cytoarchitectural composition (i.e. dentate gyrus, CA 1,3)^15,16^ For example, there is some evidence for a division of time and space in some cytoarchitectural circuits ^3^. Another possibility is that each subregion supports the same set of operations via the tri-synaptic circuit, but on different types of information received through differential connections with extrahippocampal brain regions. More anterior subregions exhibit greater functional connectivity with perirhinal cortex, while more middle and posterior regions of the hippocampus exhibit greater functional connectivity with posterior parahippocampal cortex ^35^. The perirhinal cortex is widely recognized as a region supporting complex item representations, while posterior parahippocampal cortex may support spatial and non-spatial contextual associations ^5^. A third possibility is that the differences we observed reflect more general divisions of labor that transcend the type of relation examined ^10,17^. Although we have no reason to suspect that our item-time and item-item tasks required more generalization processes (as suggested by being the only tasks associated with changes in hippocampal head), the current study cannot exclude this possibility directly. Future research is required to disentangle these possibilities.

The present research has several limitations. One potential limitation is that we did not differentiate between encoding and retrieval operations, and thus we cannot address hypotheses that anterior and posterior hippocampus preferentially support encoding and retrieval, respectively ^36^. However, it is not clear how differential support for encoding or retrieval operations could explain the structure-behavior relations we observed here, especially given identical encoding procedures, and minimization of retrieval demands using short-term memory delays. Another potential limitation is that we focused exclusively on the development of the hippocampus, while extra-hippocampal changes can additionally account for memory changes. However, the goal of this research was to examine relational memory processes in the hippocampus in a task that manipulated the type of relation. Moreover, our task used materials and procedures designed to ensure that differences in performance across relational conditions depended more strongly on hippocampally mediated associative processes ^6,8^ than on pre-frontally mediated strategic or controlled processes ^37–39^. These procedures included identical encoding procedures across relational conditions, the use of novel objects, which could not easily be labeled, and arbitrary relations among them. As discussed earlier, retrieval demands were reduced by testing memory over short delays. Finally, this research did not address how cytoarchitectural subfields in the hippocampus (i.e. dentate gyrus, CA 1-3) may account for the relations with head, body, and tail development, which should be the subject of future research and analysis.

In conclusion, we present the first evidence to establish distinct links between subregional changes in hippocampal structure to the differential development of relational memory for associations between items and space, time, and other items. These results––beyond their implication to theories of memory development—begin to disentangle the contributions of the hippocampus to three critical dimensions of relational memory.

## Materials and Methods

### Participants

Our sample included 171 participants at T1 (84 females; 143 behavioral assessments; 155 structural scans; *M*_*age*_ = 9.45 years, *SD*_*age*_ = 1.09, 7.1 – 12.0 years), 140 participants at T2 (66 females; 136 behavioral assessments, 118 structural scans; *M*_*age*_ = 10.86 years, *SD*_*age*_ = 1.22, 8.2 – 13.86 years), and 119 participants at T3 (52 females; 114 behavioral assessments, 88 structural scans; *M*_*age*_ = 12.12 years, *SD*_*age*_ = 1.31, 9.0 – 15.16 years). Item-space, item-time, and item-item memory at T1 did not significantly differ between those who returned at T2 compared to those who did not (χ^2^ = 2.61, *df* =3, *p* = .46 uncorrected), or between participants who returned for T3 and those who did not (χ^2^ = 1.31, *df* =3, *p* = .73 uncorrected). Head, body, and tail volumes did not differ at T1 in those who returned at T2 than those who did not (χ^2^s≤ 1.17, *dfs* =2, *ps* ≥ .56 uncorrected), or between participants who returned for T3 and those who did not (χ^2^s≤ 2.13, *df* s = 2, *p*s ≥ .34 uncorrected). Children were ineligible if parents reported a learning disability, neurological or psychological diagnosis requiring medication at the time of enrollment. Children were compensated for their participation. This research was conducted with the approval of the Institutional Review Board at the University of California, Davis.

### Materials and Procedures

Behavioral and imaging data were collected over two visits. The Triplet Binding Task (TBT) was administered on the first visit. Magnetic Resonance Imaging (MRI) occurred approximately one week after the behavioral assessment.

#### Triplet Binding Task

The TBT is a memory task that assesses item-time, item-space, and item-item relational memory and item-recognition memory using ^6,22^. To counter fatigue, the TBT was administered over two separate sessions on the same day. In each session, each memory type was assessed in blocks to minimize increased task-switching costs in younger children. Blocks were counterbalanced across participants. Within each assessment block, 5 encoding-retrieval phases were administered. TBT stimuli included color images of novel and obscure real-world objects unlikely to be familiar to participants; these stimuli limit the utility of semantic-based organizational memory strategies known to underlie some developmental improvements in memory ^37^.

##### Encoding Phase

Prior to each testing block, participants were instructed and tested on their understanding of the task, the relation to be encoded, and the triplet trial structure using practice encoding and retrieval phases. The encoding phase format was identical for item-time, item-space, item-item, and item-recognition encoding conditions. Each encoding phase comprised three trials. In each trial, three novel objects (i.e. triplet) were sequentially presented for one second to three locations on a computer screen, one object per location (see Figure 1B Top). A one second inter-trial fixation was then presented before proceeding to the next of the three encoding trials. To aid learning, the encoding phase was repeated a second time.

##### Retrieval Phase

Retrieval immediately followed each encoding phase. Each retrieval phase, depending on the testing block, assessed memory for item-space, item-time, or item-item relations, or item recognition memory (Figure 1B Bottom). The retrieval phase comprised three target and/or lure probes. Overall, 15 targets and 15 lures were probed in each retrieval condition.

##### Item-space

In each item-space test probe, three objects from the same encoding trial appeared together on the screen. Participants decided whether all objects appeared at their original positions or not. In target trials all objects maintain their original positions, while in lure trials the spatial positions of two objects are exchanged.

##### Item-time

In each item-time retrieval phase, three objects from the same encoding trial were sequentially presented to the center of the screen. No object appeared at their original spatial position. Participants decided whether the sequence of objects in the probe appeared in their original order or not. In target trials all objects maintain their original order, while in lure trials the ordinal position of two objects are switched.

##### Item-item

In each item-item test probe, three objects appeared on the screen at three horizontal positions. No object appeared at their original spatial position. Participants decided whether all objects had appeared together in the same trial (i.e. triplet) or not. In target trials all objects came from the same encoding trial, while in lure trials one object was exchanged with an object from another trial from the same encoding phase.

##### Item recognition

In each item-recognition test probe, three objects appeared together on the screen at three horizontal positions. No object appeared at their original spatial position. Participants decided whether all objects had previously been studied. In target trials all objects were studied, while in lure trials two of the three objects were new.

#### Magnetic Resonant Imaging

Magnetic Resonance Imaging (MRI) was acquired at the University of California, Davis Imaging Research Center in a 3T Siemens Tim Trio scanner with a 32-channel head coil. Two 7½-minute T1-weighted MPRAGE images were acquired (TE: 3.2 ms; TR: 2500 ms; in-plane resolution: 640 × 256 matrix, 0.35 mm × 0.70 mm; slice resolution: 640, 0.35 mm). Each participant’s two structural images were co-registered, averaged, and oriented so that the coronal plane was perpendicular to the long axis of the hippocampus. Each image was cropped into left and right hippocampal regions, after which retrospective bias correction was performed.

##### Hippocampal Segmentation

Hippocampal segmentation was performed using the Automatic Hippocampal Estimator using Atlas-based Delineation (AHEAD) software which implements a state-of-the-art multi-atlas joint label fusion approach to image segmentation ^40^. Briefly, manually labeled atlases of left and right hippocampus are non-linearly registered to each participant’s structural image using Advanced Normalization Tools. This produces candidate segmentations for each target’s hippocampus from which a consensus segmentation is computed using joint label fusion, an advanced weighted voting procedure ^40^. The multi-atlas of the hippocampus was produced by expert manual rater (JKL) in 14 children balanced for sex and age using an established protocol ^41^, a quantity of atlases sufficient to yield high accuracy segmentation ^42^. Each segmentation was manually reviewed for accuracy.

##### Delineation of Hippocampal Sub-Regions

Head, body, and tail subregions were delineated by blinded rater PD and JKL under an established protocol ^14^. Each subregion volume was adjusted by estimated intracranial volume (ICV) using the analysis of covariance approach ^24^. ICV estimates were obtained using previously described procedures ^15^.

### Analytical Approach

All analyses used mixed random effect models capable of accounting for within-subject dependencies in the data ^23^. Since accelerated longitudinal designs enroll participants across a range of starting ages, the effects of age comprise both the within-individual effect of age change and the between-subject effect of cross-sectional differences in age. We therefore followed the approach in which the effects of age at each time point are separated into a within-subject time-varying covariate (i.e. change in age since T1) and a between-subject time-invariant covariate (i.e. starting age at T1) ^23,25^. Given that at most only three measurement occasions were available, we did not estimate non-linear within-subject effects. However, we capitalize on the accelerated longitudinal design to test whether children of different starting ages have different within-subject trajectories. Time invariant covariates (e.g., starting age at T1) were centered at the mean of the measure at the T1. All mixed effect models included a random intercept and random slope for change in age since T1. Estimation of model parameters used restricted maximum likelihood (REML), while model comparisons used maximum likelihood (ML). Data were inspected for univariate and multivariate outliers using distribution-based outlier detection, data and Q-Q plots, Z-scoring, and Cook’s distance; outlying volume changes were identified and Winsorized at the 2^nd^ and 98^th^ percentiles. Mixed models were fitted and plotted using the lme4 (ver. 1.1), lmerTest (ver. 2.0) and effects (ver. 3.1) packages in R (ver. 3.3.1). Model comparisons were used to build up each model over baseline models, beginning with first-order effects and systematically testing inclusion of higher order interaction effects.

#### Behavioral Model

Memory scores were computed at each time point and relation as the difference between hit and false alarm rates. Models include the effects of starting age at T1, change in age, and memory relation, and control for effects of sex and item-recognition at T1. The full behavioral model is described in Table 1.

#### Hippocampal Model

We tested for main and interactive effects of starting age at T1, change in age, and hippocampal subregion, and control for effects of sex and hemisphere. The hippocampal model is described in Table 1. We also computed partial derivatives to derive the starting age at T1 in which the slope of change in age would be predicted to equal zero (i.e., the apex/base of the trajectories).

#### Brain-Behavior Model

Brain-behavior analyses examined item-time, item-space, and item-item memory separately. Each model simultaneously tested the effects of changes in hippocampal head, body, and tail on memory performance, while accounting for their volumes at T1. The brain-behavior model is described in Table 1. Model comparisons tested the effect of head, body, and tail changes together as a block, building up the model. We began by testing the change in model fit by simultaneously adding the three volume changes (as a block) over a baseline model, which included age at T1, change in age, item-recognition at T1. We then proceeded by testing the change in fit by adding the two-way interactions between changes head, body, and tail volume and change in age since T1, as a block. Likewise, the two-way interactions changes in head, body, and tail volumes with the age at T1. Lastly, we tested the change in model fit by adding the three-way interactions between changes in head, body, and tail volumes with change in age and age at T1. Finally, primary analyses summed volumes across hemispheres. Additional analyses considering left and right hippocampal structures separately were also conducted.

## Acknowledgements

Support for this research was provided by National Institute on Mental Health Research Grant (MH091109) to S.G. and S.A.B‥ J.K.L. was additionally supported by the Ruth L. Kirschstein Institutional National Research Service Award (MH073124-13 & MH073124-14).

## Declarations

The authors have no financial or non-financial competing interests to declare.

## Supplementary Information

**Figure S1 Related to Figure 4.**
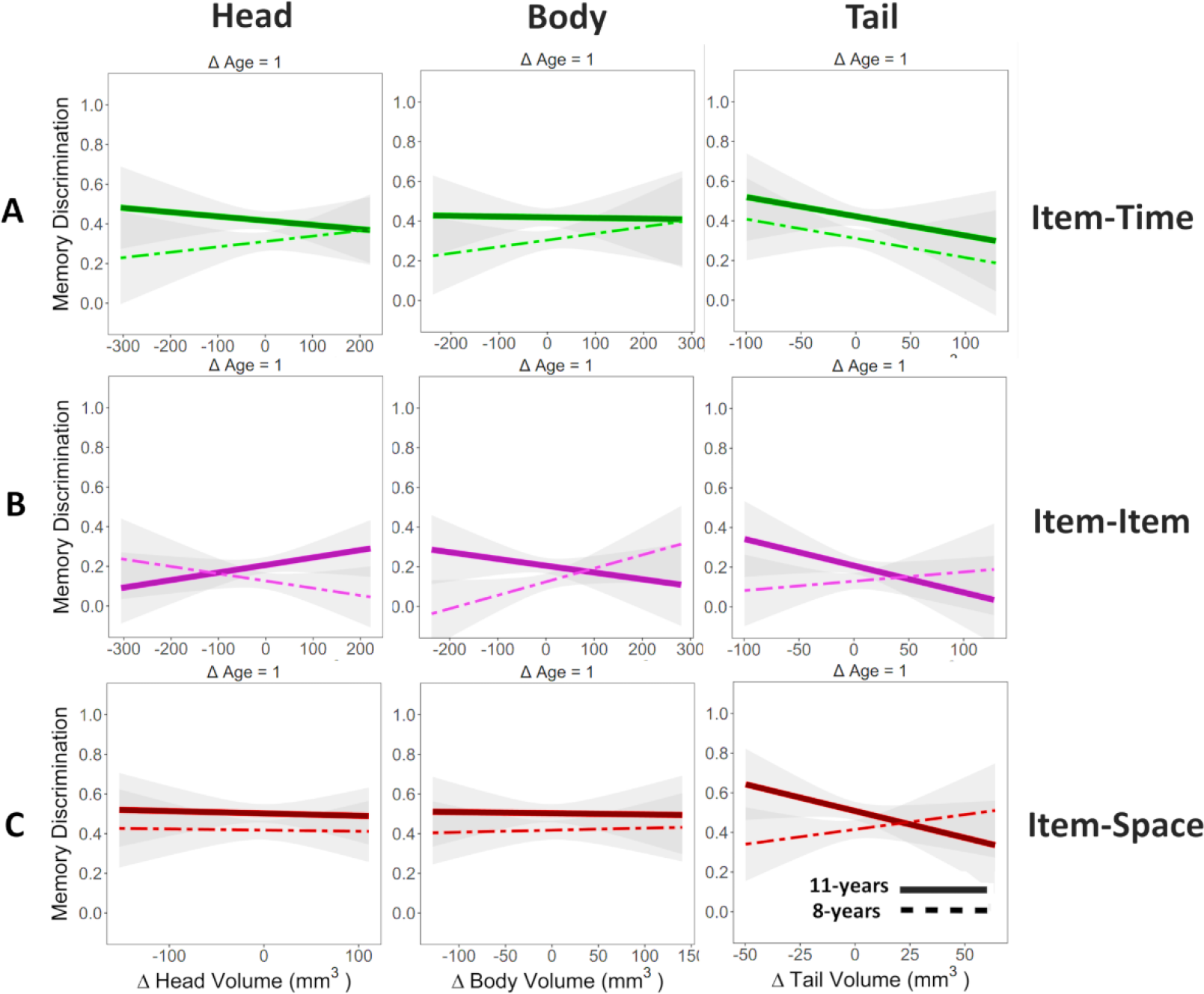
Depicting interaction between change in ICV-corrected volume and cross-sectional differences in the starting age at Time 1 at 8- and 11-years of age evaluated at a change in age since Time 1 equaling one year (ΔAge = 1). See Figure 4 for depiction of interaction after three years since Time 1; smaller changes in age corresponded to smaller differences in memory with increased sub-region ICV-corrected volume. **A.** Item-Time. **B.** Item-Item. **C.** Item-Space.

**Table S1 Related to Table 2 and Figure 2.** Relational Memory Development

| Effect | Beta | b | SE | t | p |
| --- | --- | --- | --- | --- | --- |
| (Intercept) | – | .453 | .020 | 22.9 | <.0001 |
| Item-Recognition (Mean-Centered) | .244 | .294 | .048 | 5.98 | <.0001 |
| Male | -.049 | -.026 | .021 | -1.24 | .223 |
| Start-Age (Mean-Centered) | .183 | .039 | .013 | 3.07 | .002 |
| ΔAge | .091 | .023 | .011 | 2.14 | .033 |
| Item-Time | -.218 | -.124 | .019 | -6.62 | <.0001 |
| Item-Item | -.548 | -.311 | .019 | -16.7 | <.0001 |
| Start-Age x ΔAge | -.124 | -.019 | .009 | -2.20 | .029 |
| Start-Age x Item-Time | .021 | .008 | .014 | .540 | .590 |
| Start-Age x Item-Item | .027 | -.010 | .014 | -.705 | .482 |
| ΔAge x Item-Time | .079 | .028 | .013 | 2.19 | .029 |
| ΔAge x Item-Item | .081 | .027 | .013 | 2.12 | .034 |
| Start-Age x ΔAge x Item-Time | .003 | .0007 | .011 | .066 | .950 |
| Start-Age x ΔAge x Item-Item | .002 | .0006 | .011 | .061 | .951 |

Model Fit of Fixed Effects: $\chi^2=464.3$ , $df=13$ , $p < 2.2e-16$ ; Interactions with sex were not significant, $\chi^2s \leq 4.66$ , $dfs=3$ , $ps \geq .20$ . Item-Space and female are reference categories. Thus, ΔAge and Start-Age x ΔAge represents development of Item-Space.

**Table S2 Related to Table 3.** Subregional Differences in Hippocampal Development

| **Effect** | **Beta** | **b** | **SE** | **t** | **p** |
| --- | --- | --- | --- | --- | --- |
| (Intercept) | — | 1270 | 10.0 | 127 | <.0001 |
| Male | .0008 | -2.08 | 11.7 | -.178 | .862 |
| Right Hemisphere | -.031 | 24.8 | 4.62 | 5.38 | <.0001 |
| Start-Age (Mean-Centered) | .013 | 1.41 | 6.85 | .206 | .842 |
| ΔAge | .006 | 3.20 | 3.86 | .830 | .411 |
| Hippocampal Head | -.151 | -91.3 | 7.97 | -11.5 | <.0001 |
| Hippocampal Tail | -1.02 | -1069 | 7.89 | -136 | <.0001 |
| Start-Age x ΔAge | -.031 | -9.19 | 3.64 | -2.53 | .012 |
| Start-Age x Head | -.005 | 2.66 | 7.13 | .373 | .711 |
| Start-Age x Tail | -.014 | -7.04 | 7.06 | -.997 | .322 |
| ΔAge x Head | -.022 | -12.6 | 5.27 | -2.39 | .017 |
| ΔAge x Tail | .002 | .256 | 5.23 | .049 | .960 |
| Start-Age x ΔAge x Head | .012 | 5.01 | 4.97 | 1.01 | .312 |
| Start-Age x ΔAge x Tail | .024 | 12.8 | 4.94 | 2.59 | .010 |

Model Fit of Fixed Effects: $\chi^2=6,304$ , $df=13$ , $p < 2.2e-16$ ; Interactions with hemisphere not significant: $\chi^2=4.97$ , $df=9$ , $p=.84$ . Note: Female and hippocampal body are reference categories. Thus, ΔAge and Start-Age x ΔAge represents development of the body.

**Table S3 Related to Table 3.**
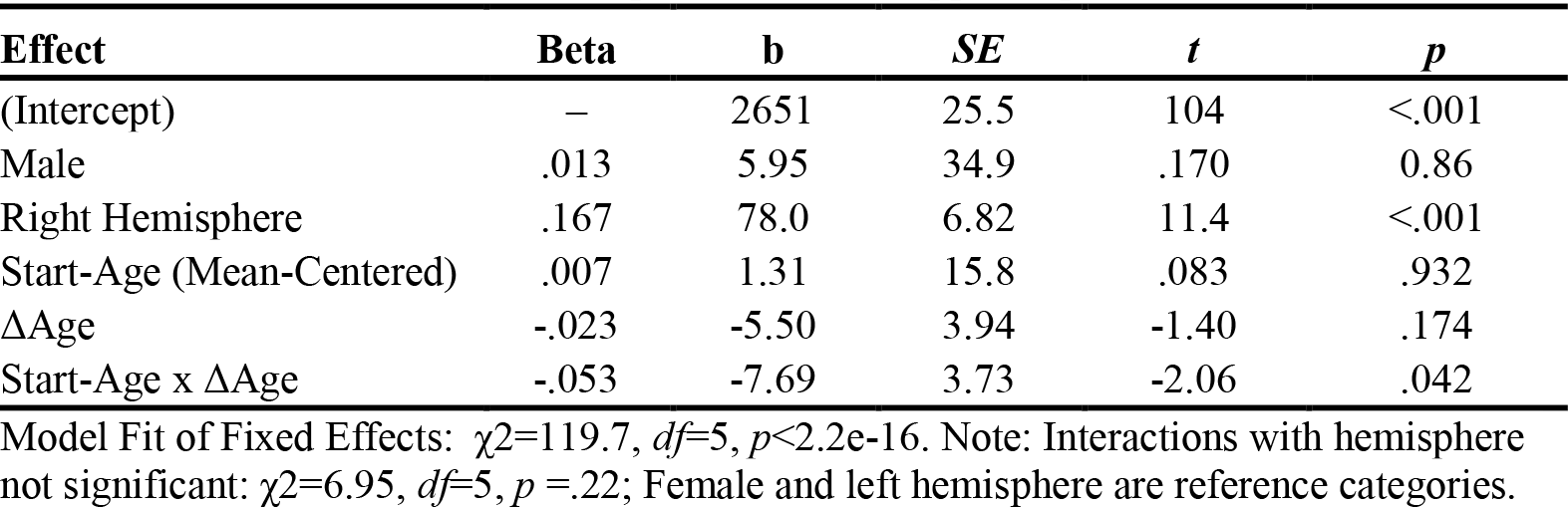
Development of Total Hippocampal Volume

**Table S4 Related to Figure 4.** Hippocampal Volume Predicting the Development of Item-Time Memory.

| Effect | Left Hippocampus |  |  |  |  | Right Hippocampus |  |  |  |  | Left and Right Hippocampal Sum |  |  |  |  |
| --- | --- | --- | --- | --- | --- | --- | --- | --- | --- | --- | --- | --- | --- | --- | --- |
|  | Beta | b | SE | t | p | Beta | b | SE | t | p | Beta | b | SE | t | p |
| (Intercept) | - | 3.1e-1 | 2.6e-2 | 12 | <.0001 | - | 3.2e-1 | 2.6e-2 | 12 | <.0001 | - | 3.2e-1 | 2.5e-2 | 13 | <.0001 |
| Item-Recognition | 0.29 | 3.3e-1 | 7.3e-2 | 4.5 | <.0001 | 0.27 | 3.1e-1 | 7.4e-2 | 4.3 | <.0001 | 0.28 | 3.2e-1 | 7.4e-2 | 4.4 | <.0001 |
| Sex [Male] | -0.041 | -2.1e-2 | 3.3e-2 | -0.65 | 0.52 | -0.061 | -3.2e-2 | 3.2e-2 | -0.99 | 0.33 | -0.053 | -2.8e-2 | 3.2e-2 | -0.86 | 0.39 |
| Start-Volume Head | -0.093 | -1.5e-4 | 1.0e-4 | -1.5 | 0.14 | -0.018 | -3.1e-5 | 1.1e-4 | -0.29 | 0.77 | -0.049 | -4.5e-5 | 5.6e-5 | -0.79 | 0.43 |
| Start-Volume Body | -0.028 | -5.6e-5 | 1.2e-4 | -0.45 | 0.65 | -0.097 | -1.8e-4 | 1.1e-4 | -1.6 | 0.12 | -0.065 | -7.0e-5 | 6.6e-5 | -1.1 | 0.29 |
| Start-Volume Tail | 0.074 | 2.2e-4 | 1.8e-4 | 1.2 | 0.24 | 0.044 | 1.3e-4 | 1.9e-4 | 0.69 | 0.49 | 0.062 | 9.8e-5 | 9.9e-5 | 0.99 | 0.32 |
| Start-Age | 0.24 | 5.6e-2 | 1.7e-2 | 3.3 | 0.0014 | 0.25 | 6.0e-2 | 1.7e-2 | 3.4 | 0.00076 | 0.25 | 5.8e-2 | 1.7e-2 | 3.4 | 0.001 |
| ΔAge | 0.25 | 6.0e-2 | 1.3e-2 | 4.7 | <.0001 | 0.27 | 6.5e-2 | 1.3e-2 | 5.2 | <.0001 | 0.26 | 6.2e-2 | 1.3e-2 | 4.7 | <.0001 |
| ΔHead | -0.11 | -4.9e-4 | 6.1e-4 | -0.81 | 0.42 | -0.12 | -6.0e-4 | 6.5e-4 | -0.92 | 0.36 | -0.063 | -1.9e-4 | 4.1e-4 | -0.47 | 0.64 |
| ΔBody | -0.2 | -9.6e-4 | 6.5e-4 | -1.5 | 0.14 | 0.02 | 9.5e-5 | 6.3e-4 | 0.15 | 0.88 | -0.056 | -1.6e-4 | 3.8e-4 | -0.43 | 0.67 |
| ΔTail | -0.25 | -2.9e-3 | 1.5e-3 | -2 | 0.05 | -0.13 | -1.6e-3 | 1.7e-3 | -0.95 | 0.34 | -0.2 | -1.4e-3 | 9.5e-4 | -1.5 | 0.14 |
| Start-Age x ΔAge | -0.092 | -1.5e-2 | 1.2e-2 | -1.3 | 0.21 | -0.15 | -2.4e-2 | 1.2e-2 | -2 | 0.044 | -0.13 | -2.1e-2 | 1.2e-2 | -1.7 | 0.095 |
| Start-Age x ΔHead | -0.34 | -1.4e-3 | 5.5e-4 | -2.5 | 0.014 | -0.22 | -9.5e-4 | 5.5e-4 | -1.7 | 0.084 | -0.26 | -6.4e-4 | 3.2e-4 | -2 | 0.048 |
| Start-Age x ΔBody | -0.3 | -1.4e-3 | 6.6e-4 | -2.1 | 0.035 | -0.22 | -9.4e-4 | 5.8e-4 | -1.6 | 0.11 | -0.22 | -6.3e-4 | 3.7e-4 | -1.7 | 0.096 |
| Start-Age x ΔTail | -0.001 | -1.1e-5 | 1.4e-3 | -0.008 | 0.99 | -0.078 | -9.5e-4 | 1.8e-3 | -0.54 | 0.59 | -0.037 | -2.6e-4 | 1.0e-3 | -0.25 | 0.8 |
| ΔAge x ΔHead | 0.12 | 2.8e-4 | 3.1e-4 | 0.89 | 0.37 | 0.18 | 4.6e-4 | 3.6e-4 | 1.3 | 0.2 | 0.072 | 1.1e-4 | 2.2e-4 | 0.5 | 0.62 |
| ΔAge x ΔBody | 0.31 | 7.0e-4 | 3.3e-4 | 2.1 | 0.035 | 0.004 | 9.1e-6 | 3.1e-4 | 0.029 | 0.98 | 0.14 | 2.0e-4 | 1.9e-4 | 1.1 | 0.29 |
| ΔAge x ΔTail | 0.15 | 8.7e-4 | 7.9e-4 | 1.1 | 0.27 | 0.11 | 6.4e-4 | 9.0e-4 | 0.71 | 0.48 | 0.11 | 4.0e-4 | 5.0e-4 | 0.8 | 0.42 |
| Start-Age x ΔAge x ΔHead | 0.4 | 8.4e-4 | 2.9e-4 | 2.9 | 0.0042 | 0.29 | 6.4e-4 | 3.1e-4 | 2.1 | 0.038 | 0.33 | 4.1e-4 | 1.9e-4 | 2.2 | 0.027 |
| Start-Age x ΔAge x ΔBody | 0.35 | 8.4e-4 | 3.5e-4 | 2.4 | 0.018 | 0.31 | 7.0e-4 | 3.2e-4 | 2.2 | 0.028 | 0.29 | 4.2e-4 | 1.9e-4 | 2.2 | 0.032 |
| Start-Age x ΔAge x ΔTail | 0.032 | 1.8e-4 | 7.3e-4 | 0.25 | 0.81 | 0.22 | 1.4e-3 | 9.4e-4 | 1.5 | 0.15 | 0.12 | 4.2e-4 | 5.5e-4 | 0.77 | 0.44 |

Note: Female is reference sex. For unstandardized betas, volume is in cubic millimeters and age is in years.

**Table S5 Related to Figure 4.** Hippocampal Volume Predicting the Development of Item-Item Memory.

| Effect | Left Hippocampus |  |  |  |  | Right Hippocampus |  |  |  |  | Left and Right Hippocampal Sum |  |  |  |  |
| --- | --- | --- | --- | --- | --- | --- | --- | --- | --- | --- | --- | --- | --- | --- | --- |
|  | Beta | b | SE | t | p | Beta | b | SE | t | p | Beta | b | SE | t | p |
| (Intercept) | - | 1.2e-1 | 2.3e-2 | 5.3 | <.0001 | - | 1.2e-1 | 2.3e-2 | 5.2 | <.0001 | - | 1.2e-1 | 2.2e-2 | 5.4 | <.0001 |
| Item-Recognition | 0.15 | 1.4e-1 | 6.1e-2 | 2.4 | 0.019 | 0.15 | 1.4e-1 | 6.2e-2 | 2.3 | 0.024 | 0.16 | 1.5e-1 | 6.2e-2 | 2.5 | 0.013 |
| Sex [Male] | -0.015 | -6.6e-3 | 2.7e-2 | -0.24 | 0.81 | -8.2e-05 | -3.6e-5 | 2.7e-2 | -0.001 | 0.99 | -0.0048 | -2.1e-3 | 2.7e-2 | -0.077 | 0.94 |
| Start-Volume Head | -0.018 | -2.5e-5 | 8.6e-5 | -0.29 | 0.77 | -0.011 | -1.6e-5 | 8.9e-5 | -0.18 | 0.85 | -0.013 | -1.0e-5 | 4.7e-5 | -0.21 | 0.83 |
| Start-Volume Body | -0.012 | -2.1e-5 | 1.0e-4 | -0.2 | 0.84 | 0.025 | 3.9e-5 | 9.7e-5 | 0.4 | 0.69 | 0.025 | 2.3e-5 | 5.6e-5 | 0.41 | 0.69 |
| Start-Volume Tail | 0.034 | 8.3e-5 | 1.5e-4 | 0.55 | 0.59 | 0.0012 | 2.9e-6 | 1.6e-4 | 0.019 | 0.99 | 0.018 | 2.4e-5 | 8.2e-5 | 0.29 | 0.78 |
| Start-Age | 0.21 | 4.2e-2 | 1.5e-2 | 2.7 | 0.0068 | 0.2 | 3.8e-2 | 1.5e-2 | 2.5 | 0.014 | 0.2 | 3.9e-2 | 1.5e-2 | 2.6 | 0.012 |
| ΔAge | 0.25 | 5.1e-2 | 1.1e-2 | 4.7 | <.0001 | 0.27 | 5.5e-2 | 1.1e-2 | 5.2 | <.0001 | 0.27 | 5.5e-2 | 1.1e-2 | 5 | <.0001 |
| ΔHead | 0.026 | 1.0e-4 | 5.4e-4 | 0.19 | 0.85 | -0.13 | -5.5e-4 | 5.8e-4 | -0.94 | 0.35 | -0.048 | -1.2e-4 | 3.5e-4 | -0.35 | 0.73 |
| ΔBody | -0.19 | -7.4e-4 | 5.8e-4 | -1.3 | 0.2 | 0.1 | 4.1e-4 | 5.5e-4 | 0.74 | 0.46 | -0.00071 | -1.7e-6 | 3.4e-4 | -0.005 | 1 |
| ΔTail | -0.068 | -6.6e-4 | 1.3e-3 | -0.5 | 0.62 | -0.14 | -1.4e-3 | 1.5e-3 | -0.95 | 0.34 | -0.15 | -8.7e-4 | 8.5e-4 | -1 | 0.31 |
| Start-Age x ΔAge | -0.11 | -1.5e-2 | 1.0e-2 | -1.5 | 0.15 | -0.11 | -1.5e-2 | 1.0e-2 | -1.4 | 0.16 | -0.081 | -1.1e-2 | 1.0e-2 | -1.1 | 0.29 |
| Start-Age x ΔHead | 0.075 | 2.5e-4 | 2.0e-4 | 1.3 | 0.2 | 0.062 | 2.2e-4 | 2.0e-4 | 1.1 | 0.28 | 0.12 | 2.5e-4 | 1.2e-4 | 2.1 | 0.039 |
| Start-Age x ΔBody | -0.12 | -4.9e-4 | 2.2e-4 | -2.2 | 0.029 | -0.06 | -2.2e-4 | 2.2e-4 | -1 | 0.32 | -0.13 | -3.0e-4 | 1.4e-4 | -2.2 | 0.028 |
| Start-Age x ΔTail | 0.01 | 9.0e-5 | 5.0e-4 | 0.18 | 0.86 | 0.034 | 3.5e-4 | 5.8e-4 | 0.6 | 0.55 | 0.015 | 8.5e-5 | 3.3e-4 | 0.26 | 0.8 |
| ΔAge x ΔHead | 0.1 | 1.9e-4 | 2.7e-4 | 0.71 | 0.48 | 0.25 | 5.2e-4 | 3.0e-4 | 1.7 | 0.087 | 0.17 | 2.1e-4 | 1.8e-4 | 1.1 | 0.26 |
| ΔAge x ΔBody | 0.34 | 6.5e-4 | 2.8e-4 | 2.3 | 0.022 | -0.027 | -5.1e-5 | 2.6e-4 | -0.2 | 0.85 | 0.1 | 1.2e-4 | 1.6e-4 | 0.76 | 0.45 |
| ΔAge x ΔTail | 0.068 | 3.4e-4 | 6.7e-4 | 0.51 | 0.61 | 0.11 | 5.4e-4 | 7.6e-4 | 0.72 | 0.48 | 0.13 | 3.8e-4 | 4.3e-4 | 0.88 | 0.38 |

Note: Female is reference sex. For unstandardized betas, volume is in cubic millimeters and age is in years.

**Table S6 Related to Figure 4.** Hippocampal Volume Predicting the Development of Item-Space Memory.

| Effect | Left Hippocampus |  |  |  |  | Right Hippocampus |  |  |  |  | Left and Right Hippocampal Sum |  |  |  |  |
| --- | --- | --- | --- | --- | --- | --- | --- | --- | --- | --- | --- | --- | --- | --- | --- |
|  | Beta | b | SE | t | p | Beta | b | SE | t | p | Beta | b | SE | t | p |
| (Intercept) | - | 0.46 | 0.026 | 18 | <.0001 | - | 0.46 | 0.025 | 18 | <.0001 | - | 0.46 | 0.025 | 18 | <.0001 |
| Item-Recognition | 0.31 | 3.5e-1 | 7.1e-2 | 4.9 | <.0001 | 0.31 | 3.5e-1 | 7.2e-2 | 4.8 | <.0001 | 0.31 | 3.4e-1 | 7.2e-2 | 4.7 | <.0001 |
| Sex [Male] | -0.089 | -4.5e-2 | 3.2e-2 | -1.4 | 0.16 | -0.083 | -4.2e-2 | 3.2e-2 | -1.3 | 0.19 | -0.08 | -4.0e-2 | 3.2e-2 | -1.3 | 0.21 |
| Start-Volume Head | -0.00091 | -1.4e-6 | 1.0e-4 | -0.014 | 0.99 | -0.027 | -4.6e-5 | 1.0e-4 | -0.45 | 0.66 | -0.012 | -1.1e-5 | 5.6e-5 | -0.2 | 0.84 |
| Start-Volume Body | -0.097 | -1.9e-4 | 1.2e-4 | -1.6 | 0.12 | -0.042 | -7.6e-5 | 1.1e-4 | -0.68 | 0.5 | -0.068 | -7.2e-5 | 6.5e-5 | -1.1 | 0.27 |
| Start-Volume Tail | 0.0045 | 1.3e-5 | 1.8e-4 | 0.071 | 0.94 | 0.039 | 1.1e-4 | 1.8e-4 | 0.61 | 0.54 | 0.019 | 2.9e-5 | 9.8e-5 | 0.3 | 0.76 |
| Start-Age | 0.27 | 6.1e-2 | 1.7e-2 | 3.6 | 0.0004 | 0.28 | 6.3e-2 | 1.7e-2 | 3.8 | 0.0003 | 0.27 | 6.2e-2 | 1.7e-2 | 3.7 | 0.0003 |
| ΔAge | 0.1 | 2.4e-2 | 1.3e-2 | 1.8 | 0.071 | 0.12 | 2.9e-2 | 1.3e-2 | 2.2 | 0.028 | 0.1 | 2.4e-2 | 1.4e-2 | 1.8 | 0.078 |
| ΔHead | 0.055 | 2.5e-4 | 6.1e-4 | 0.41 | 0.69 | -0.086 | -4.2e-4 | 6.5e-4 | -0.65 | 0.52 | 0.012 | 3.6e-5 | 4.1e-4 | 0.087 | 0.93 |
| ΔBody | -0.0065 | -3.0e-5 | 6.5e-4 | -0.046 | 0.96 | -0.018 | -8.3e-5 | 6.3e-4 | -0.13 | 0.9 | 0.0087 | 2.5e-5 | 3.9e-4 | 0.065 | 0.95 |
| ΔTail | -0.1 | -1.2e-3 | 1.5e-3 | -0.78 | 0.44 | -0.14 | -1.7e-3 | 1.7e-3 | -0.99 | 0.32 | -0.13 | -8.7e-4 | 9.7e-4 | -0.9 | 0.37 |
| Start-Age x ΔAge | -0.17 | -2.7e-2 | 1.2e-2 | -2.2 | 0.027 | -0.21 | -3.4e-2 | 1.2e-2 | -2.7 | 0.0077 | -0.20 | -3.1e-2 | 1.3e-2 | -2.5 | 0.014 |
| Start-Age x ΔHead | -0.062 | -2.4e-4 | 5.5e-4 | -0.44 | 0.66 | -0.055 | -2.3e-4 | 5.5e-4 | -0.42 | 0.68 | 0.0019 | 4.6e-6 | 3.3e-4 | 0.014 | 0.99 |
| Start-Age x ΔBody | -0.2 | -9.2e-4 | 6.7e-4 | -1.4 | 0.17 | -0.11 | -4.7e-4 | 5.8e-4 | -0.81 | 0.42 | -0.13 | -3.7e-4 | 3.8e-4 | -0.97 | 0.33 |
| Start-Age x ΔTail | 0.18 | 1.9e-3 | 1.4e-3 | 1.4 | 0.16 | -0.33 | -4.0e-3 | 1.8e-3 | -2.3 | 0.025 | -0.012 | -8.2e-5 | 1.0e-3 | -0.078 | 0.94 |
| ΔAge x ΔHead | -0.035 | -7.7e-5 | 3.2e-4 | -0.25 | 0.81 | 0.13 | 3.2e-4 | 3.6e-4 | 0.88 | 0.38 | -0.015 | -2.2e-5 | 2.3e-4 | -0.096 | 0.92 |
| ΔAge x ΔBody | 0.093 | 2.1e-4 | 3.3e-4 | 0.62 | 0.54 | 0.025 | 5.5e-5 | 3.2e-4 | 0.17 | 0.86 | 0.06 | 8.3e-5 | 1.9e-4 | 0.43 | 0.67 |
| ΔAge x ΔTail | 0.0033 | 1.9e-5 | 8.0e-4 | 0.024 | 0.98 | 0.16 | 9.6e-4 | 9.2e-4 | 1 | 0.3 | 0.071 | 2.5e-4 | 5.1e-4 | 0.49 | 0.63 |
| Start-Age x ΔAge x ΔHead | 0.091 | 1.9e-4 | 2.9e-4 | 0.63 | 0.53 | 0.12 | 2.4e-4 | 3.1e-4 | 0.78 | 0.44 | 0.034 | 4.2e-5 | 1.9e-4 | 0.22 | 0.82 |
| Start-Age x ΔAge x ΔBody | 0.17 | 3.9e-4 | 3.5e-4 | 1.1 | 0.27 | 0.2 | 4.4e-4 | 3.2e-4 | 1.4 | 0.18 | 0.18 | 2.5e-4 | 2.0e-4 | 1.3 | 0.21 |
| Start-Age x ΔAge x ΔTail | -0.19 | -1.0e-3 | 7.4e-4 | -1.4 | 0.18 | 0.41 | 2.4e-3 | 9.6e-4 | 2.5 | 0.012 | 0.024 | 8.3e-5 | 5.6e-4 | 0.15 | 0.88 |

Note: Female is reference sex. For unstandardized betas, volume is in cubic millimeters and age is in years.

